# CCR2 recruits monocytes to the lung, while CX3CR1 modulates positioning of monocyte-derived CD11c^pos^ cells in the lymph node during pulmonary tuberculosis

**DOI:** 10.1101/2025.02.07.637199

**Authors:** Alexander Mohapatra, Zachary Howard, Joel D. Ernst

## Abstract

Infection by *Mycobacterium tuberculosis* (Mtb) continues to cause more than 1 million deaths annually, due to pathogen persistence in lung macrophages and dendritic cells derived from blood monocytes. While accumulation of monocyte-derived cells in the Mtb-infected lung partially depends on the chemokine receptor CCR2, the other chemoattractant receptors regulating trafficking remain undefined. We used mice expressing knock-in/knockout reporter alleles of *Ccr2* and *Cx3cr1* to interrogate their expression and function in monocyte-derived populations of the lungs and draining mediastinal lymph nodes during Mtb infection. CCR2 and CX3CR1 expression varied across monocyte-derived subsets stratified by cell surface Ly6C expression in both organs. We found that expression of CCR2 predicted dependence of monocyte-derived cells on the receptor for lung and lymph node accumulation. CCR2-deficient mice were also observed to have worsened lung and lymph node Mtb burden. While CX3CR1 deficiency, alone or in combination with CCR2 deficiency, did not affect cell frequencies or lung Mtb control, its absence was associated with altered positioning of monocyte-derived dendritic cells in mediastinal lymph nodes. We found that combined loss of *Ccr2* and *Cx3cr1* also worsened Mtb control in the mediastinal lymph node, suggesting a rationale for the persistent expression of CX3CR1 among monocyte-derived cells in pulmonary tuberculosis.

**IMPORTANCE:** *Mycobacterium tuberculosis* is the respiratory pathogen responsible for the deadliest infectious disease worldwide. Susceptible humans exhibit ineffective immune responses, in which infected phagocytes are not able to eliminate the pathogen. Since recruited monocyte-derived cells serve as reservoirs for persistent infection, understanding how these phagocytes accumulate in the lung and why they are unable to eliminate Mtb can inform development of therapies that can synergize with antimicrobials to achieve faster and more durable Mtb elimination. Monocyte-derived cells express the chemokine receptors CCR2 and CX3CR1, but the role of the latter in Mtb infection remains poorly defined. The significance of our study is in elucidating the roles of these receptors in the trafficking of monocyte-derived cells in the infected lung and mediastinal lymph node. These data shed light on the host response in tuberculosis and in other pulmonary infections.

## INTRODUCTION

Tuberculosis remains one of the deadliest infectious diseases worldwide, causing more than 1 million deaths annually [1]. Humans who progress to disease following inhalation of *M. tuberculosis* (Mtb) exhibit ineffective immune responses, in which infected phagocytes are not able to eliminate the mycobacteria [2]. Since recruited monocyte-derived cells serve as reservoirs for persistent infection [3, 4, 5, 6], understanding how these phagocytes accumulate in the lung and why they are unable to eliminate Mtb can inform development of therapies that can synergize with antimicrobials to achieve faster and more durable Mtb elimination.

Following phagocytosis by alveolar macrophages (AMs) [7, 8, 9], Mtb establishes a durable niche in the lung parenchyma within phagocytes derived from blood monocytes [3, 4, 5, 6, 10]. Monocyte-derived macrophages become the dominant infected cells within the lungs of mice, while monocyte-derived dendritic cells (DCs) traffic Mtb to draining lymph nodes (LNs) for T cell activation [9, 11, 12, 13, 14]. However, T cells do not effectively eliminate Mtb from monocyte-derived lung macrophages [4, 5, 15, 16, 17]. Moreover, blood monocytes are continuously recruited to the infected lung [18, 19], creating a renewable reservoir of Mtb. Therefore, a key question is how monocytes traffic to the lungs and LNs during Mtb infection.

Two phenotypes of blood monocytes, classical and non-classical, are defined by expression of the chemokine receptors CCR2 and CX3CR1, respectively, suggesting their importance to monocyte biology [20]. CCR2 deficiency impairs monocyte egress from the bone marrow and may affect recruitment to sites of inflammation [21, 22, 23]. Cell surface components of Mtb [24] and Type I interferons [25] induce the CCR2 ligand CCL2 in Mtb infection, enhancing monocyte recruitment to the lung. Studies of infected, CCR2-deficient mice demonstrate that the receptor is essential for maximal accumulation of monocyte-derived cells in the lungs [26, 27]. Diminished monocyte recruitment is associated with impaired Mtb control, likely due to reduced activation of Mtb-specific T cells [13, 26, 27, 28, 29]. However, mononuclear phagocyte recruitment is not completely abrogated in the absence of CCR2, indicating that one or more other chemoattractant receptors contributes to this response to infection. Expression of the CX3CR1 ligand CX3CL1 has not been characterized in Mtb infection, but it is highly expressed in the human lung at baseline [30]. CX3CR1^pos^ monocyte-derived cells have been observed to phagocytose *M. bovis* bacille Calmette-Guérin during pulmonary infection [31]. However, Mtb-infected *Cx3cr1^-/-^* mice exhibited similar mycobacterial control to that of wildtype mice [32], although phagocyte accumulation in the lungs was not reported in that study. Given that monocytes highly express CCR2 and CX3CR1 and that monocyte trafficking to the Mtb-infected lung is not completely abrogated in CCR2-deficient mice, we generated mice deficient in both CCR2 and CX3CR1 and used them to evaluate accumulation of monocyte-derived cells and Mtb control in the lungs and mediastinal LNs (MLNs) following aerosol infection.

We used the knock-in/knockout *Ccr2^RFP/RFP^* [33] and *Cx3cr1^GFP/GFP^* [34] strains to breed double knockout (DKO) mice and double heterozygous mice, for tracking *Ccr2*- and *Cx3cr1*-expressing phagocytes during pulmonary Mtb infection. Mice that are heterozygous for these alleles exhibit normal migratory properties attributable to the remaining intact copy of each chemokine receptor gene and express the respective reporter. Mice that are homozygous at each targeted locus are null for the respective chemokine receptor. We observed variable receptor expression among monocyte-derived subsets and differential dependence on these receptors for accumulation of these cells in the lung parenchyma and the MLN. While CCR2-deficient and DKO mice had similar impairment of lung Mtb control, compared with that in CCR2-replete mice, combined absence of Ccr2 and Cx3cr1 was associated with altered positioning of monocyte-derived DCs in the MLN and with higher MLN Mtb burdens.

## RESULTS

### Variable expression of Ccr2 and Cx3cr1 by mononuclear phagocyte subsets in *M. tuberculosis*-infected mice

Since *Ccr2* and *Cx3cr1* are genetically linked on *M. musculus* chromosome 9, studying the effects of dual receptor deficiency on phagocyte trafficking required mating of F1 littermates until a crossing-over event generated a dual reporter allele (Fig. 1A). To study expression of these receptors in specific cell subsets, we infected *Ccr2^RFP/+^*; *Cx3cr1^GFP/+^* (denoted “Het” hereafter) mice with Mtb and used flow cytometry to analyze cells isolated from lungs and MLNs 4 weeks post-infection (Fig. 1B). We measured RFP and GFP median fluorescence intensity (MFI), as reflective of receptor gene expression, in monocyte-derived macrophages, monocyte-derived DCs, AMs, and neutrophils (gating in Supplemental Fig. 1). Expression was also assessed in CD103^pos^ conventional DCs, which are rarely infected in the lung but present Mtb-derived antigens in MLNs [13, 14].

**FIG 1.**
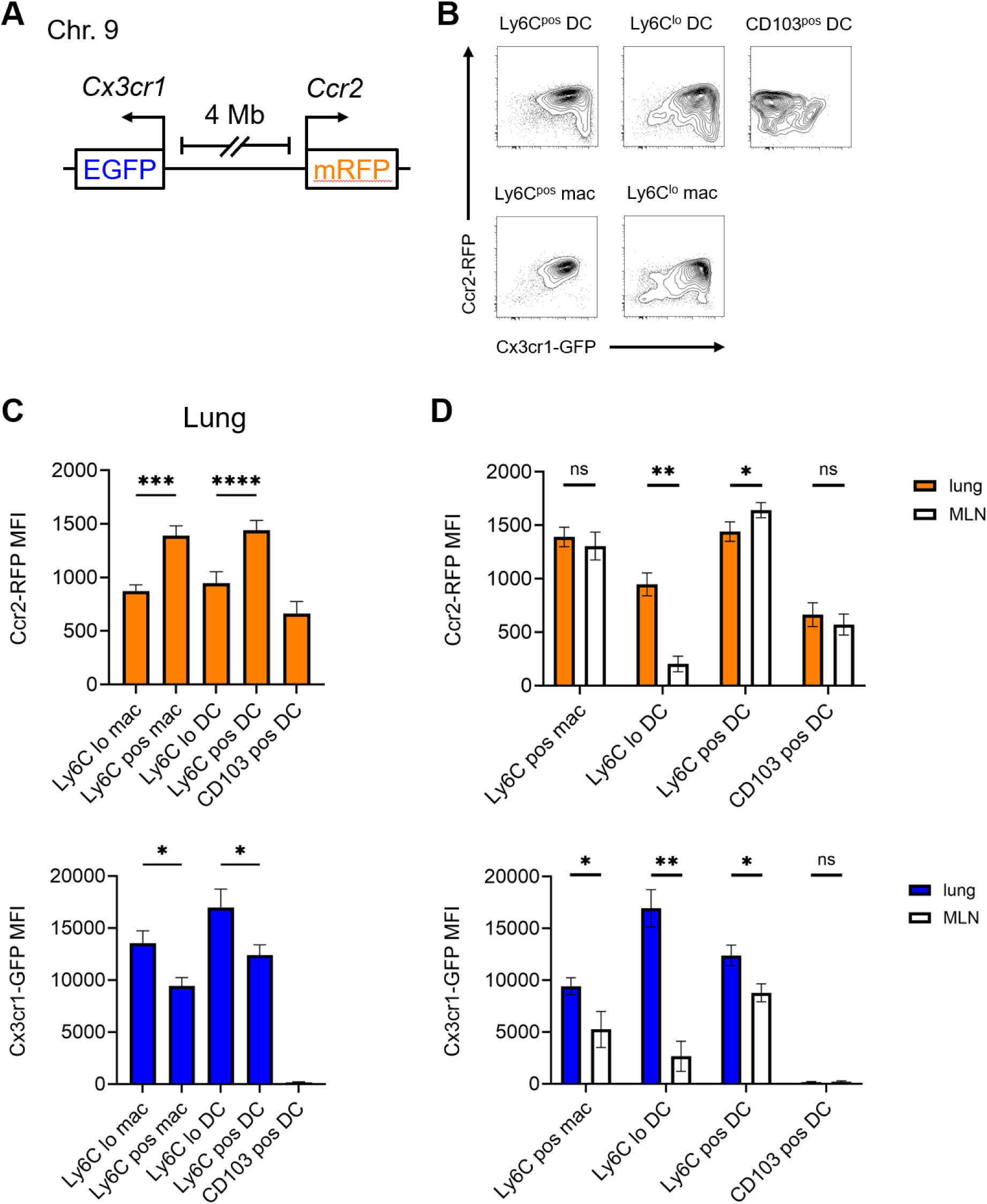
A dual *Ccr2*-*Cx3cr1* reporter mouse demonstrates varied receptor expression by cell type and location. (A) Schematic of the dual reporter genes on chromosome 9. (B-D) *Ccr2^RFP/+^*; *Cx3cr1^GFP/+^* mice were infected with Mtb, then euthanized at 4 weeks post-infection for analysis of lung and MLN cells by flow cytometry (gating in Supplemental Fig. 1). (B) Representative contour plots of the indicated populations from the lungs of infected mice (“mac”: monocyte-derived macrophage). (C) Median fluorescence intensities (MFIs) of RFP (top) and GFP (bottom) in lung cells. (D) Comparison of RFP (top) and GFP (bottom) MFIs among equivalent lung and MLN populations (Ly6C^lo^ macrophages were not observed in significant numbers in the MLN). Data in (B-D) are representative of at least 2 experiments with 4 mice per genotype. All results presented as arithmetic mean + SD. *, p<0.05; **, p<0.005; ***, p<0.0005; ****, p<0.00005; ns, not significant; by one-way ANOVA with multiple comparisons (C) or *t* test (D).

We assessed RFP and GFP expression by organ and cell type (Fig. 1B). Intravascular (IV) anti-CD45 antibody injection prior to euthanasia was used to distinguish intravascular populations (CD45-IV^pos^), including Ly6C^hi^ CCR2^hi^ CX3CR1^lo^ classical and Ly6C^lo^ CCR2^lo^ CX3CR1^hi^ non-classical monocytes, from those in the lung parenchyma (CD45-IV^neg^). In Mtb-infected mice, we have previously used adoptive transfer of monocytes to determine that parenchymal populations are derived from circulating classical monocytes, which transition from Ly6C^pos^ to Ly6C^lo^ over time [19] and assume macrophage-like or DC-like phenotypes in the lung [12]. We predicted that these monocyte-derived cells would have similar chemokine receptor expression kinetics as do blood monocytes, which lose CCR2 and Ly6C expression and acquire CX3CR1 expression when transitioning from classical monocytes into non-classical monocytes [35]. This prediction was correct, as Ly6C^lo^ cells had lower RFP (*Ccr2* reporter) and higher GFP (*Cx3cr1* reporter) expression than did Ly6C^pos^ populations (Fig. 1C). Parenchymal CD103^pos^ DCs did not express significant levels of GFP but had detectable RFP expression. AMs and neutrophils expressed minimal RFP or GFP (Supplemental Fig. 2), as expected [21, 34, 36, 37].

We previously found that both Ly6C^pos^ and Ly6C^lo^ monocyte-derived populations migrate from the lung parenchyma to the MLN in Mtb-infected mice [19], so we predicted that expression of CCR2 and CX3CR1 would be similar between lung and MLN populations. While MLN Ly6C^pos^ macrophages and DCs resembled lung populations in RFP and GFP MFI, MLN Ly6C^lo^ DCs had significantly lower expression of both reporters than did equivalent cells in the lung (Fig. 1D). CD103^pos^ DCs in the MLN had expression of CCR2 and CX3CR1 comparable to that in lung cells. For all lung and MLN populations evaluated, RFP and GFP expression peaked at 4 weeks post-infection and were significantly decreased by 7 weeks (Supplemental Fig. 3A-B).

### Lung and MLN phagocytes differ in their dependence on CCR2 and CX3CR1

Our chemokine receptor expression data suggested that Ly6C^lo^ monocyte-derived cells might utilize CX3CR1 for recruitment to the lung parenchyma, potentially explaining the partial trafficking defect observed with CCR2 deficiency alone. To test this hypothesis, we measured accumulation of monocyte-derived cells in the lungs and MLNs of Mtb-infected mice deficient in CCR2, CX3CR1, or both receptors. We confirmed that Mtb-infected Het mice had similar accumulation of lung phagocyte populations as did wildtype (*Ccr2^+/+^*; *Cx3cr1^+/+^*) mice (Supplemental Fig. 4A) and could be used as controls. In the lung at 4 weeks post-infection, accumulation of Ly6C^pos^ macrophages and DCs declined by more than 80 percent in CCR2-deficient (*Ccr2^RFP/RFP^*) and DKO mice, while CX3CR1-deficient (*Cx3cr1^GFP/GFP^*) mice had preserved trafficking relative to Het mice (Fig. 2A). Accumulation of Ly6C^lo^ macrophages and DCs, however, was less dependent on CCR2, although additional loss of CX3CR1 did not further reduce the frequencies of these populations. Whereas Ly6C^pos^ DCs were the most prevalent monocyte-derived subset in Het and CX3CR1-deficient mice, the residual monocyte-derived cells in CCR2-deficient and DKO mice were Ly6C^lo^ DCs (Fig. 2B). Overall, our data suggest that while CCR2 expression predicts accumulation of monocyte-derived cells in the lungs of Mtb-infected mice, CX3CR1 expression does not. Conversely, despite expressing detectable RFP (Fig. 1C), accumulation of CD103^pos^ DCs was not dependent on CCR2 (Fig. 2A). Absence of CCR2 was also associated with increased parenchymal neutrophils. Given that neutrophils did not express CCR2, as indicated by RFP expression (Supplemental Fig. 2) and in the literature [21], this is likely an indirect effect potentially due to higher mycobacterial burdens (Fig. 4A).

**FIG 2.**
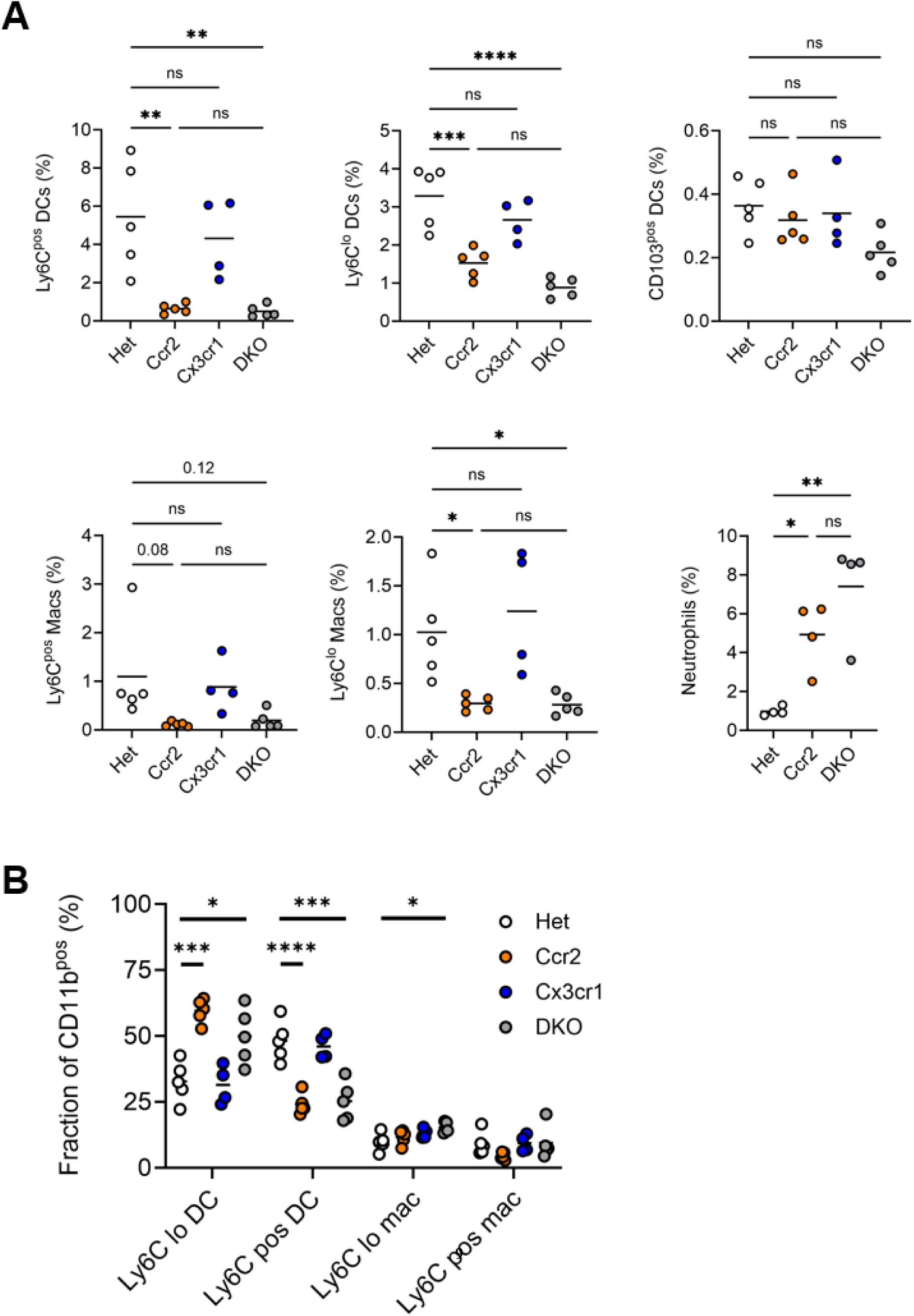
Phagocyte frequencies in the lungs of Mtb-infected CCR2-deficient, CX3CR1-deficient, and double knockout mice. *Ccr2^RFP/+^*; *Cx3cr1^GFP/+^* (Het), *Ccr2^RFP/RFP^* (Ccr2), *Cx3cr1^GFP/GFP^* (Cx3cr1), and DKO mice were infected with Mtb for 4 weeks, and lungs were harvested for analysis. (A) Frequencies of the indicated lung populations among live cells by flow cytometry. (B) Proportions of the indicated populations among CD11b^pos^ cells. Data in (A-B) are representative of at least 2 experiments with at least 4 mice per genotype. Horizontal bar represents the arithmetic mean in all graphs. Significance assessed by one-way ANOVA with multiple comparisons.

The significantly lower CCR2 and CX3CR1 expression in MLN Ly6C^lo^ DCs, relative to lung Ly6C^lo^ DCs, suggested that trafficking of this population to the MLN does not depend on these receptors. This was confirmed, as accumulation of Ly6C^lo^ and CD103^pos^ DCs was similar in the MLNs of infected Het, single, and double knockout mice (Fig. 3A). MLN Ly6C^pos^ macrophages and DCs, which expressed the CCR2 reporter, were nearly absent in CCR2-deficient and DKO mice. Consequently, Ly6C^lo^ DCs were the dominant monocyte-derived population in CCR2-deficient and DKO MLNs (Fig. 3B). Neutrophils were the only MLN phagocyte significantly affected by CX3CR1 deficiency. Absence of both CCR2 and CX3CR1 was associated with an increase in neutrophil frequency (Fig. 3A), again presumed to be an indirect effect (Supplemental Fig. 2).

**FIG 3.**
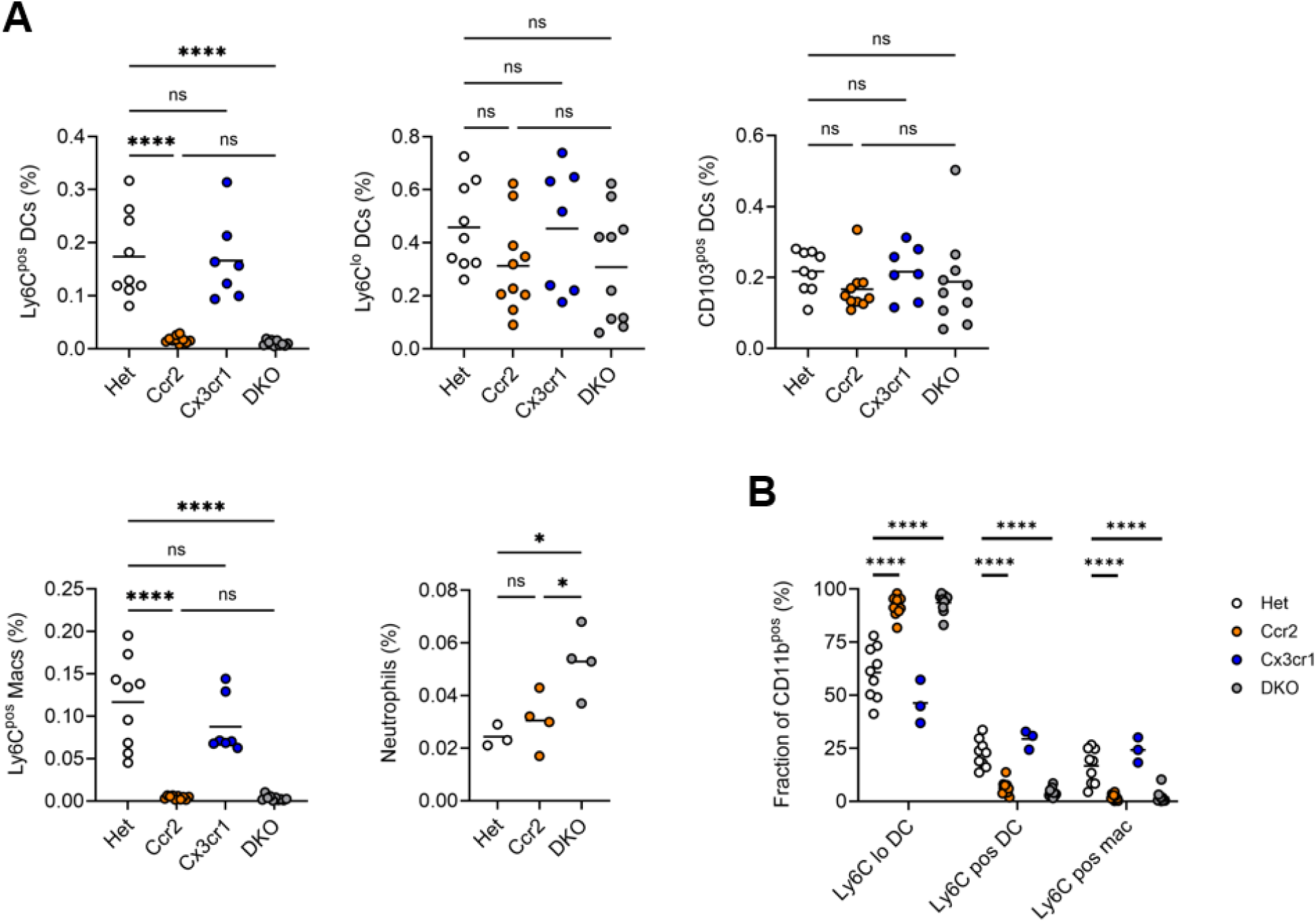
Phagocyte frequencies in the MLNs of Mtb-infected CCR2-deficient, CX3CR1-deficient, and double knockout mice. Mice were infected, as in Fig. 2, and MLNs were harvested for analysis. (A) Frequencies of the indicated MLN populations among live cells by flow cytometry. (B) Proportions of the indicated populations among CD11b^pos^ cells. Data in (A-B) are representative of at least 2 experiments with at least 4 mice per genotype. Horizontal bar represents the arithmetic mean in all graphs. Significance assessed by one-way ANOVA with multiple comparisons.

### Mtb-infected cells exhibit altered positioning in MLNs of DKO mice

To further examine the potential effects of chemokine receptor deficiency on the recruitment and function of mononuclear cells in the MLN, we infected CCR2-deficient and DKO mice with an Mtb strain expressing yellow fluorescent protein (YFP) [38]. This permitted identification of infected phagocytes in the lungs and MLNs with simultaneous detection of the GFP and RFP reporters (Supplemental Fig. 1). To measure transport of Mtb to the MLN by infected lung cells, we quantitated the frequency of CCR7^pos^ phagocytes at 4 weeks post-infection and observed that CCR2 deficiency was associated with more CCR7^pos^ YFP^pos^ Ly6C^lo^ DCs in the lungs (Fig. 4A). However, additional loss of CX3CR1 did not further increase the frequency of these cells. Some CCR7^pos^ YFP^pos^ CD11b^lo^ conventional DCs (both CD103^lo^ and CD103^pos^; not shown) were observed in the lungs, but these were less prevalent than CD11b^pos^ Ly6C^lo^ monocyte-derived DCs and were not affected by absence of CCR2 or CX3CR1. We next quantitated Mtb-infected cells in the MLN and observed that YFP^pos^ DCs could not be detected in Het mice (not shown). However, YFP^pos^ CD11b^lo^ and Ly6C^lo^ DCs were observed in the MLNs of CCR2-deficient and DKO mice. Animals lacking both CCR2 and CX3CR1 had fewer YFP^pos^ CD11b^lo^ DCs but a similar frequency of YFP^pos^ Ly6C^lo^ DCs, compared with mice deficient in CCR2 alone (Fig. 4B). Although the frequency of Ly6C^lo^ DCs in the MLNs of infected mice was not affected by deficiency of CCR2 or CX3CR1 (Fig. 3A), we asked whether Ly6C^lo^ DCs in DKO mice had altered positioning within MLNs. We used immunofluorescence microscopy of MLN sections to quantitate CD11c^pos^ cells in non-follicular regions (defined as B220^neg^; Fig. 4C), including T cell zones (Supplemental Fig. 5A). Whereas CD11c^pos^ cells in CCR2-deficient mice tended to be adjacent to follicular zones, CD11c^pos^ cells were further away from B220^pos^ areas in DKO mice. We did not directly compare localization of CD11c^pos^ cells in Het mice with that in CCR2-deficient or DKO mice because of the disparity in Ly6C^pos^ DCs (Fig. 3A) and Mtb burden (Fig. 5B). Since we observed more neutrophils in the MLNs of infected DKO mice (Fig. 3A), we also analyzed co-localization of Ly6G and YFP in MLN sections and found few infected neutrophils (Supplemental Fig. 5B).

**FIG 4.**
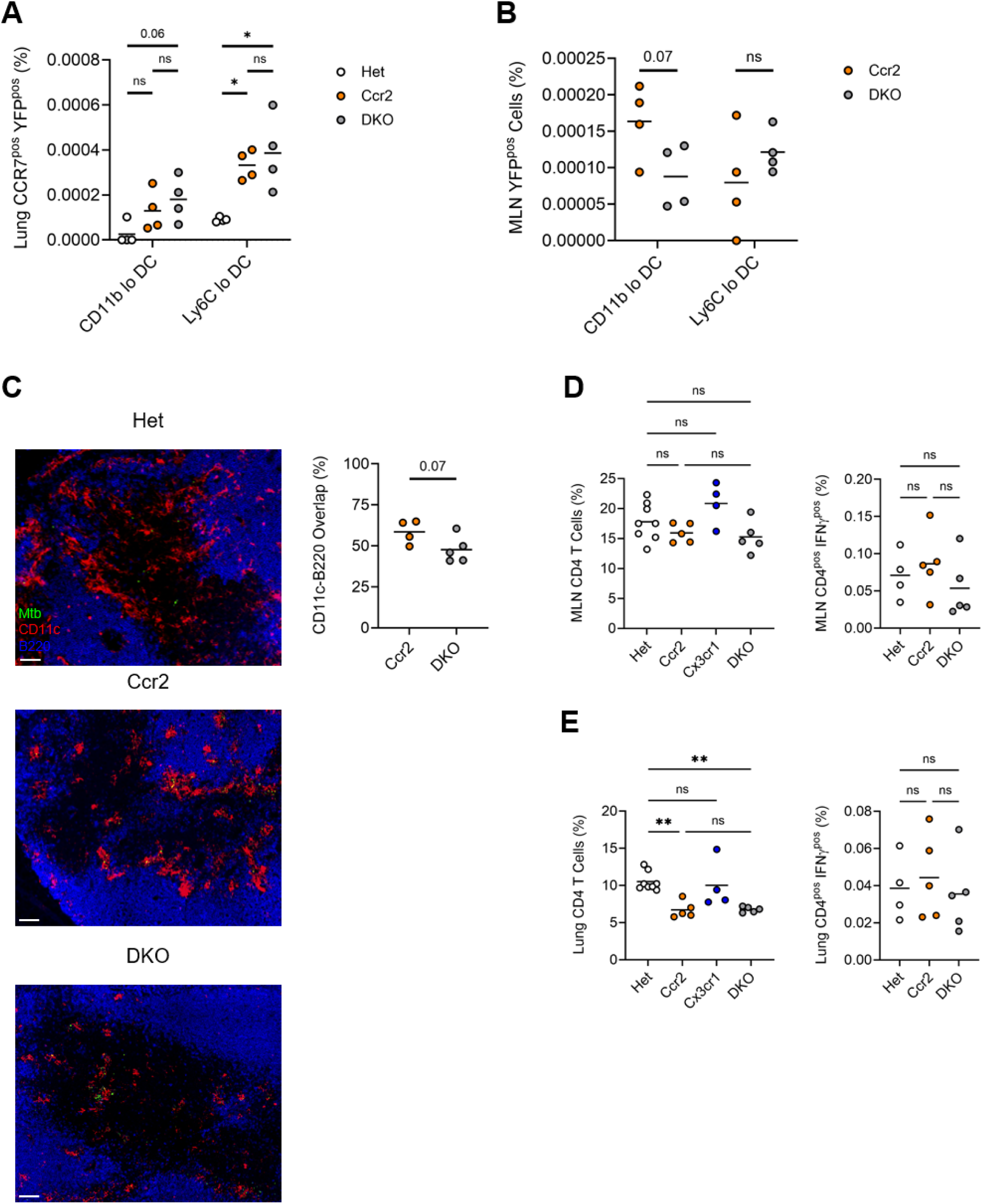
Mtb-infected cells exhibit altered positioning in the MLNs of double knockout mice. Lungs and MLNs were harvested from mice infected with YFP-expressing Mtb for 4 weeks. (A) Frequency of CCR7^pos^ YFP^pos^ cells of the indicated subsets among total lung cells. (B) Frequency of YFP^pos^ cells of the indicated subsets among total MLN cells. (C) Representative immunofluorescence staining of MLN sections from Het, Ccr2, or DKO mice. Scale bar is 70 *µ*m. Graph shows the percentage of CD11c-positive staining that overlaps with B220-positive staining in non-follicular zones, compiled from 3 experiments. (D-E) Frequencies of total and IFN*γ*^pos^ CD4 T cells in the MLN (D) and lung (E). Data in (A-B, D-E) are representative of 2 experiments with 4 mice per genotype. Horizontal bar represents the arithmetic mean in all graphs. Significance assessed by one-way ANOVA with multiple comparisons (A, D-E) or *t* test (B-C).

**FIG 5.**
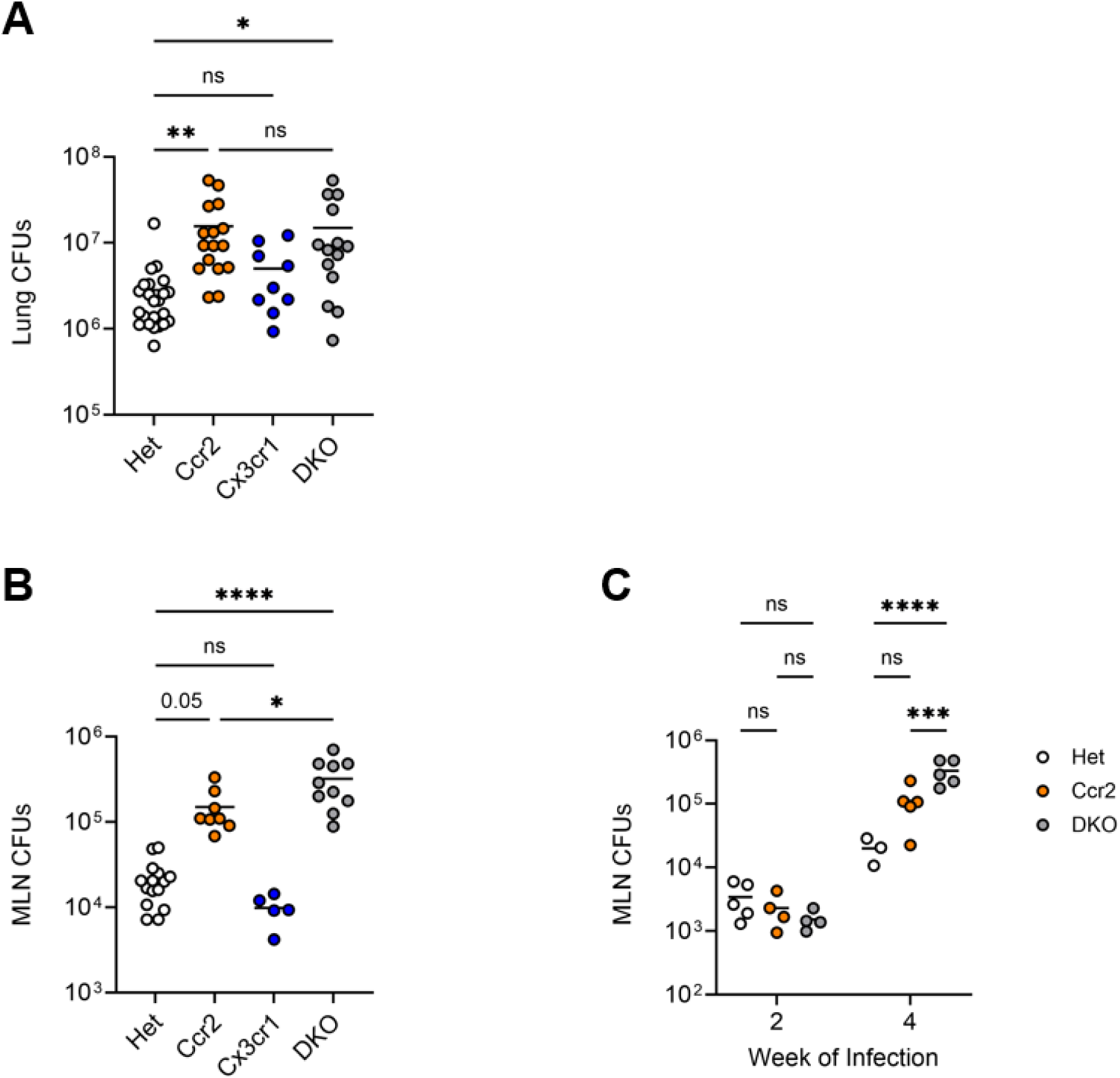
Double knockout mice have greater mycobacterial burden in MLNs than CCR2-deficient mice. Mice were infected for 2 or 4 weeks, then lungs and MLNs were harvested. (A) Quantitation of Mtb bacilli in the lungs of infected mice by colony formation (CFUs) on agar. Data are compiled from 4 experiments. (B) Quantitation of Mtb bacilli in the MLNs of mice infected for 4 weeks. Data are compiled from 3 experiments. (C) MLN CFUs after 2 and 4 weeks of infection. Data are representative of 2 experiments with at least 3 mice per genotype. Horizontal bar represents the arithmetic mean in all graphs. Significance assessed by one-way (A-B) or two-way (C) ANOVA with multiple comparisons.

We then assessed how the change in MLN DC populations induced by CCR2 deficiency affected accumulation and polarization of CD4 T cells. In the MLN, the frequency of total CD4 T cells was not affected by the absence of CCR2 or CX3CR1 (Fig. 4D). In contrast, CD4 T cell frequency in the lung parenchyma was reduced by 35 percent in CCR2-deficient mice, when compared with Het mice (Fig. 4E). There was no difference in lung CD4 T cell frequency between mice lacking CCR2 alone and those lacking both receptors. While CCR2 deficiency was associated with reduced overall frequency of CD4 T cells in the lungs, the frequency of IFN*γ*^pos^ CD4 T cells in both the MLN (Fig. 4D) and the lung (Fig. 4E) was not affected by single or dual deficiency of CCR2 or CX3CR1.

### Combined deficiency of CCR2 and CX3CR1 worsens Mtb control in the MLN but not the lung compared with CCR2 deficiency alone

We next quantitated viable Mtb bacilli in the lungs of Het, Ccr2 and Cx3cr1 single knockouts, and DKO mice at 4 weeks post-infection. We found significantly more Mtb colony-forming units (CFUs) in the lungs of CCR2-deficient and DKO mice (Fig. 5A), compared to those in Het mice (which had similar Mtb burdens as wildtype *Ccr2^+/+^*; *Cx3cr1^+/+^* mice; Supplemental Fig. 4C). Mtb CFUs in mice lacking both receptors did not differ from those in mice lacking CCR2 alone. In contrast, mycobacterial burdens did not differ in the lungs of CX3CR1-deficient mice compared with those in Het mice. Together, these results with CCR2-deficient mice are similar to those in prior studies [26, 27, 29] and reveal that deficiency of CX3CR1 does not impact lung bacterial burdens, even when combined with CCR2 deficiency.

In contrast to the findings in the lungs, in the MLNs of mice infected with Mtb for 4 weeks, deficiency of both CCR2 and CX3CR1 was associated with higher mycobacterial burdens compared to those in single knockouts (Fig. 5B). As observed in the lungs, CCR2-deficient mice had more bacilli in MLNs than did CX3CR1-deficient mice, which were similar to those in Het mice. To determine whether the difference in bacilli observed in the MLNs at 4 weeks post-infection was present at an earlier phase of the infection, we quantified Mtb bacilli at 2 weeks post-infection. MLN Mtb burden was lower than at 4 weeks post-infection, as expected, but was not affected by CCR2 or CX3CR1 deficiency at the earlier time point (Fig. 5C).

## DISCUSSION

We generated and used dual reporter and knockout mice to define the expression and roles of the chemokine receptors CCR2 and CX3CR1 in lung mononuclear phagocytes during Mtb infection. Distinct monocyte-derived cell subsets were observed to have specific patterns of *Ccr2* and *Cx3cr1* expression related to cell surface Ly6C and localization within the lung parenchyma or the MLN. We confirmed that CCR2-deficient mice have significantly reduced accumulation of Ly6C^pos^ monocyte-derived cells in the lung and MLN, with higher lung and MLN Mtb burden. CX3CR1 deficiency, alone or in combination with CCR2 deficiency, did not affect monocyte-derived populations or Mtb control in the lung but was associated with higher mycobacterial burdens in the MLN and altered positioning of monocyte-derived Ly6C^lo^ DCs.

Dual reporter mice revealed dynamic regulation of *Ccr2* and *Cx3cr1* in monocyte-derived cells following extravasation into the Mtb-infected lung. Our finding that Ly6C expression directly correlates with that of *Ccr2* and inversely correlates with *Cx3cr1* among lung monocyte-derived cells, independent of whether these cells adopted a macrophage-like or a DC-like phenotype, mimics the distinction between Ly6C^hi^ CCR2^hi^ CX3CR1^lo^ and Ly6C^lo^ CCR2^lo^ CX3CR1^hi^ monocytes in the blood [20]. The cells we profiled are extravascular, and we have previously shown that Ly6C^hi^ bone marrow monocytes differentiate into Ly6C^lo^ cells over time within the lung parenchyma [19], suggesting that they are not derived from blood Ly6C^lo^ cells. Therefore, our results are consistent with a shared program between blood and lung monocytes, whereby these cells repress *Ccr2* and induce *Cx3cr1* expression, possibly via induction of *N4ra1* [39]. Similar receptor expression kinetics have been observed among monocyte-derived cells in the peritoneum [34], skin [40, 41], skeletal muscle [42], myocardium [43], gut [44], and liver [45]. Our finding that CCR2 deficiency results in a greater proportion of Ly6C^lo^ DCs in the infected lung implies that these cells are a more stable, terminal population, occupying a niche that still fills, but at a slower rate, when accumulation of Ly6C^pos^ monocytes is reduced. Accumulation of a long-lived monocyte-derived population may underlie our observation that both *Ccr2* and *Cx3cr1* expression appear to decline in all phagocytes analyzed as the infection progresses, though we cannot exclude that systemic changes in monocyte receptor expression also occur with prolonged infection.

We and others have previously shown that CCR2 [13, 26, 28] and CCR7 [10, 11, 19, 46] contribute to accumulation of monocyte-derived DCs and to T cell activation in the MLNs of Mtb-infected mice. Our data on *Ccr2* expression and its role in trafficking of monocyte-derived populations to the lung and MLN suggest a model in which CCR2 facilitates monocyte accumulation in the infected lung, consistent with other studies [21, 22, 23], while CCR7 regulates MLN entry. Since our data and those from other studies [26, 27, 28, 29] suggest that monocyte trafficking to the Mtb-infected lung is delayed, rather than completely abrogated, in CCR2-deficient mice, other chemokine receptors must contribute to monocyte trafficking. However, we also do not observe a role for CX3CR1 in accumulation of monocyte-derived cells in the Mtb-infected lung or MLN, despite detectable expression. The LN result agrees with other data showing that CX3CR1^pos^ monocyte-derived DCs traffic to LN T cell zones in the spleen [34] and the intestines [47] in a CX3CR1-independent manner.

Our observation that DKO mice have increased MLN Mtb bacilli compared to single knockouts and controls reveals a novel role for CX3CR1. We [10, 14, 17, 19] and others [13] have previously shown that viable Mtb are transported into the MLN from the lung 1-2 weeks post-infection via monocyte-derived cells, which subsequently transfer Mtb antigens, but not live mycobacteria, to conventional DCs for T cell priming. Since we did not observe an increase in MLN Mtb burden in DKO mice until 4 weeks post-infection and since neither receptor is required for MLN entry, the worsened Mtb control is unlikely to be explained by trafficking of more infected cells to the MLN from the lung. Rather, we found that altered positioning of Ly6C^lo^ DCs away from follicular regions, due to CX3CR1 deficiency, was associated with Mtb survival within these cells. This phenotype is likely only apparent in DKO mice, and at a later stage of infection, because of two effects. First, significantly reduced trafficking of Ly6C^pos^ DCs to the MLN impairs T cell activation, resulting in higher lung Mtb burden and inflammation (as indicated by increased neutrophil accumulation) over time. Second, delayed accumulation of Ly6C^lo^ DCs in the lung eventually provides an alternative path for Mtb to reach the MLN. Conversely, Cx3cr1 single knockout mice have an earlier influx of Ly6C^pos^ DCs to the MLN, resulting in antigen transfer to conventional DCs that do not express the receptor and normal activation of Mtb-specific T cells.

While CX3CR1^pos^ monocyte-derived macrophages in the uninfected lung have been associated with bronchiole-associated nerves [48, 49], the tissue distribution of the chemokine itself has not been described, so the functional role of the receptor for these macrophages remains unclear. Similarly, expression of CX3CR1 ligands has not been defined in MLNs. However, our data suggest that a chemokine gradient does exist in Mtb infection and directs Ly6C^lo^ DCs to the T cell zone-B cell follicle border. In the adaptive immune response to Mtb, it may not be impactful because we have previously shown that infected Ly6C^lo^ DCs poorly present antigen to T cells [14, 17]. It is unclear why altered positioning of Ly6C^lo^ DCs affects Mtb survival or antigen export to conventional DCs. Our data also establish that CX3CR1 does not act redundantly with CCR2 to traffic monocytes to the infected lung.

Several groups have studied Mtb infection of Ccr2 [27, 29] or Cx3cr1 [32] single knockout mice. Our data are consistent with these prior studies but further extend them by demonstrating a unique role for CX3CR1 in the MLN that may explain its coordinated regulation with CCR2 in monocyte-derived cells at sites of inflammation. More research is needed to fully define the regulation of monocyte trafficking to the Mtb-infected lung and MLN and reveal opportunities for eliminating Mtb-infected phagocytes.

## MATERIALS AND METHODS

### Mice

C57BL/6 mice (8–12 weeks old) were obtained from Jackson Laboratory. Mice infected with Mtb were housed in the Animal Biosafety Level 3 facility. Ccr2-RFP and Cx3cr1-GFP mice were purchased from The Jackson Laboratory then bred to generate *Ccr2^RFP/RFP^*; *Cx3cr1^GFP/GFP^*. All animal protocols used here were approved by the University of California, San Francisco, Animal Care and Use Committee.

### Mycobacterial strains, growth, and aerosol infection

Mtb H37Rv transformed with pMV261 containing the gene expressing superfolder yellow fluorescent protein was obtained from the laboratory of Christopher Sassetti [38]. Mycobacteria were grown in Middlebrook 7H9 medium (BD) supplemented with 10% (v/v) ADC (albumin, dextrose, catalase), 0.05% Tween 80, 0.2% glycerol and 110 *µ*g/ml hygromycin. Mice were infected with H37Rv or H37Rv::pMV261-sfYFP via aerosol using an inhalation exposure unit from Glas-Col, as previously described [4, 19]. Mid-log cultures were centrifuged at 800 x *g* to pellet clumps. Clump-free cultures were then diluted, and 5 mL inoculum was added to the nebulizer. Target dose was 100 CFUs/mouse. Infection dose was determined by plating lung homogenates 24 hours post-infection on 7H11 agar plates and enumerating CFUs after 3 weeks incubation at 37°C.

### Tissue harvests and processing for flow cytometry

Mice were anesthetized by inhalation of isoflurane and then retro-orbitally injected with 1 *µ*g of anti-CD45 antibody (clone 30-F11) in 150 *µ*L of PBS. Three minutes after antibody injection mice were euthanized by CO_2_ inhalation and cervical dislocation. Lungs and MLNs were collected and placed in HBSS containing 50 *µ*g/mL Liberase TM (Sigma), 30 *µ*g/mL DNase I (Sigma) or RPMI-1640 with 5% HI-FBS (v/v), respectively. Lymph nodes were mashed through 70-*µ*m cell strainers. Lungs were incubated at 37°C for 30 minutes, then processed with a gentleMACS dissociator (Miltenyi) and mashed through 70-*µ*m strainers. Aliquots for plating on 7H11 agar for CFU enumeration were taken prior to centrifugation. Following centrifugation, lung cell suspensions were dissolved in ACK lysis buffer (Gibco) for red blood cell lysis. Lung and MLN single-cell suspensions were washed and resuspended in PBS with 3% HI-FBS (v/v), 2 mM EDTA (FACS buffer).

For staining of surface antigens, cells were washed with PBS, then stained with viability dye Live-or-Dye 750/777 (Biotium) and anti-CD16/32 (clone 2.4G2; Fc receptor blockade) for 15 min at 4°C. Cells were then washed and resuspended in FACS buffer with fluorochrome-conjugated anti-CCR7 antibody (clone 4B12) for 30 minutes at 37°C. After washing, cells were resuspended in FACS buffer with a cocktail of fluorochrome-conjugated antibodies against CD3 (17A2), CD4 (GK1.5), CD11b (M1/70), CD11c (N418), CD19 (6D5), CD90.2 (30-H12), CD103 (2E7), Ly6C (HK1.4), Ly6G (1A8), MHCII I-A and I-E (M5/114.15.2), NK1.1 (S17016D), and Siglec F (S17007L) for 25 minutes at 4°C. Cells were then washed with PBS, fixed with 4% paraformaldehyde (v/v) for 30 minutes at room temperature, and washed again in PBS.

For staining of intracellular IFN*γ*, cells were fixed with BD Cytofix/Cytoperm solution following staining of surface antigens, then washed in BD Perm/Wash buffer. After washing, cells were resuspended in BD Perm/Wash buffer with anti-IFN*γ* antibodies (XMG1.2) for 30 minutes at 4°C. Afterward, cells were washed in Perm/Wash buffer, then in PBS. Samples were acquired using an LSR II (BD) conventional flow cytometer or an Aurora spectral flow cytometer (Cytek Biosciences). See Supplementary Fig. 1 for gating.

### Lymph node immunofluorescence

Mice were euthanized, as above, and intact MLNs were embedded in OCT, then rapidly frozen in 100% ethanol with dry ice. Blocks were cut into 10-*µ*m sections by cryostat (Leica), and sections were transferred to slides and crosslinked using a CryoJane tape transfer system. Slides were fixed in 4% paraformaldehyde overnight, then stored in PBS at 4°C until immunofluorescence staining. For staining, sections were blocked with 10% goat serum for 30 minutes, then stained with antibodies against CD11c and B220 (clone RA3-6B2) for 1 hour. After washing, sections were stained with isotype-specific, fluorochrome-conjugated secondary antibodies, followed by DAPI. Sections were mounted with VECTASHIELD Vibrance Antifade Medium (Vector Laboratories). Images were captured using the 20X objective of a Nikon Ti inverted microscope with a DS-Qi2 camera. Frames in an 8 x 8 grid were stitched together and deconvoluted using NIS Elements software (Nikon).

At least 3 MLN sections 50 *µ*m apart were analyzed per mouse. For each MLN image, B220-negative regions were manually selected in ImageJ (National Institutes of Health). Masks of channels corresponding to B220 or CD11c were then generated using manually adjusted thresholds and a 20-*µ*m border (to capture cell interactions 1-2 cell lengths apart) for each B220-negative region. Overlap, defined as the amount of CD11c signal adjacent to B220-positive regions, was calculated as the area common to both the B220 and CD11c masks (using “AND” operator) as a fraction of the total CD11c mask area. The total pixel area of each B220-negative region was used to generate a weighted average of overlap for the MLN sections from each mouse.

### Data analysis

Flow cytometry data were analyzed using SpectroFlo 3.3 (Cytek) and FlowJo 10.10.0. Median fluorescence intensities for GFP and RFP were calculated after subtracting values recorded for equivalent populations from Mtb-infected *Ccr2^+/+^*; *Cx3cr1^+/+^* mice. GraphPad Prism 10.4.1 (GraphPad) was used for graphical presentation and statistical analysis.

## ACKNOWLEDGMENTS

We thank Brian Norris for early contributions to this work and Lucas Chen for technical assistance. The UCSF Division of Experimental Medicine Core Immunology Lab and Parnassus Flow CoLab (RRID: *SCR 018206*) facilitated generation of flow cytometry data. We also acknowledge the UCSF Center for Advanced Light Microscopy for assistance with image acquisition. This research was supported by NIH grants R01 AI051242 (J.D.E), U01 AI166309 (J.D.E), T32 HL007185 (A.M.), F32 HL162424 (A.M.), and P30 AI168440/R25 AI147375 (A.M.). The funders had no role in study design, data collection and interpretation, or the decision to submit the work for publication.

A.M. and J.D.E. conceived the research. A.M. and Z.H. performed the experiments. A.M. analyzed the data. A.M. and J.D.E. wrote the manuscript. J.D.E. acquired funding and supervised the project.

---

**Supplemental Figure 1:**
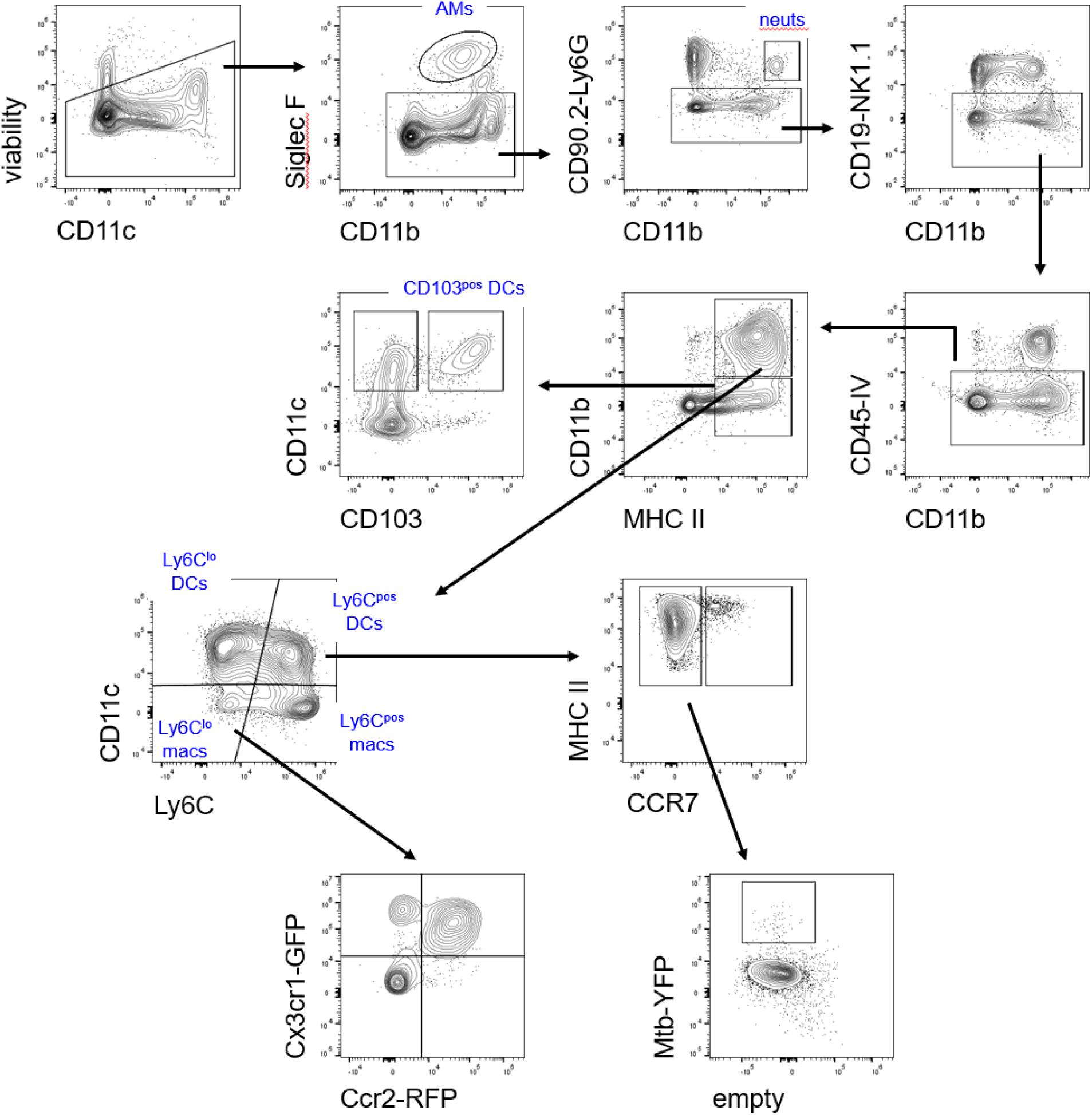
Gating strategy for lung and MLN populations. Representative contour plots from the lungs of Mtb-infected mice (“neuts”: neutrophils). Gating for MLN samples was similar. For analysis of T cells, CD3^pos^ CD4^pos^ events were gated from live cells (not shown).

**Supplemental Figure 2:**
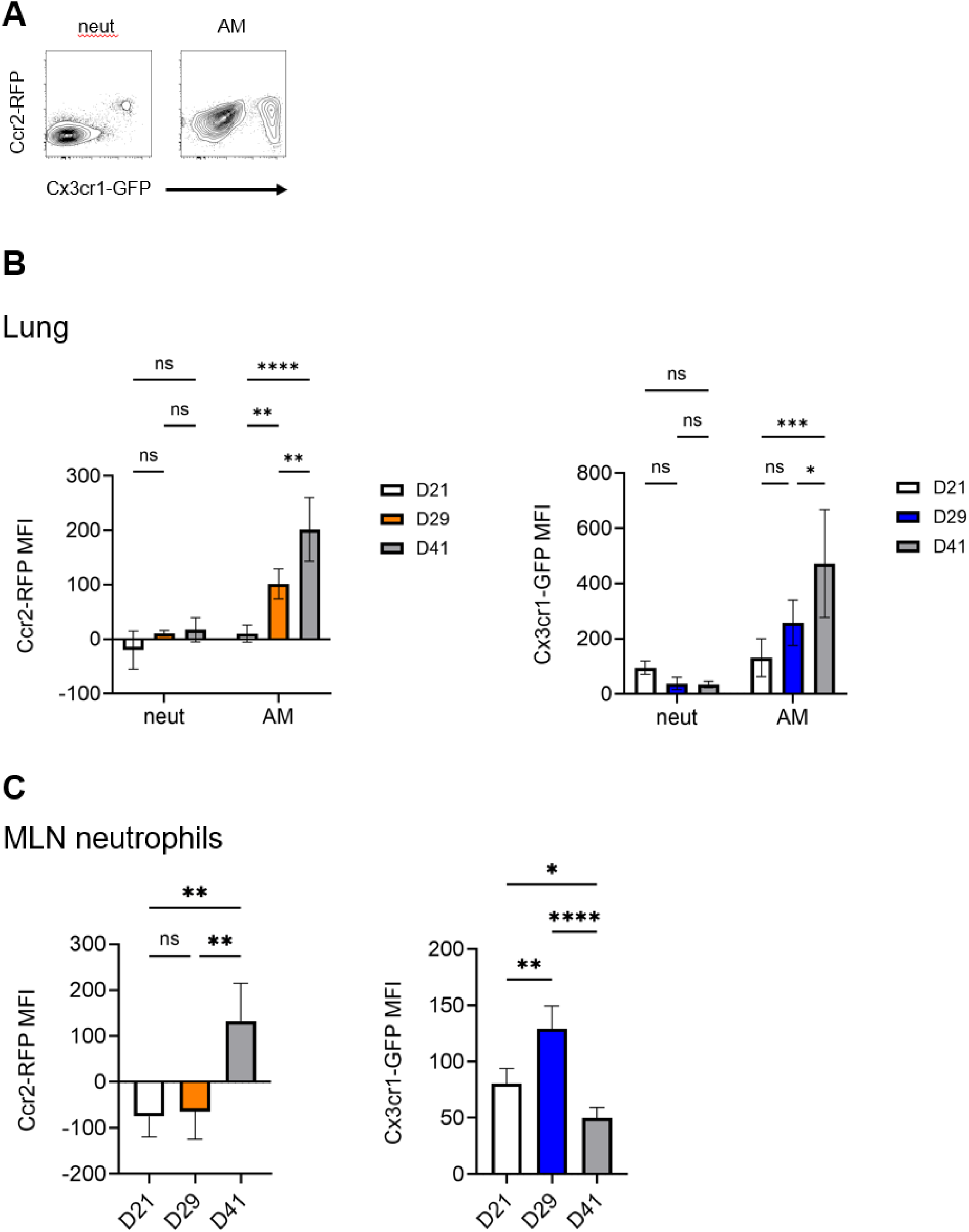
*Ccr2* and *Cx3cr1* expression in alveolar macrophages and neutrophils. (A) Representative contour plots of neutrophils and AMs from the lungs of mice infected with Mtb for 4 weeks. (B-C) *Ccr2^RFP/+^*; *Cx3cr1^GFP/+^* mice were infected with Mtb for the indicated times, then MFIs of RFP and GFP in lung neutrophils and AMs (B) and MLN neutrophils (C) were measured by flow cytometry. All results presented as mean + SD and are derived from 1 experiment with 4 mice per genotype. Significance assessed by two-way (B) or one-way (C) ANOVA with multiple comparisons.

**Supplemental Figure 3:**
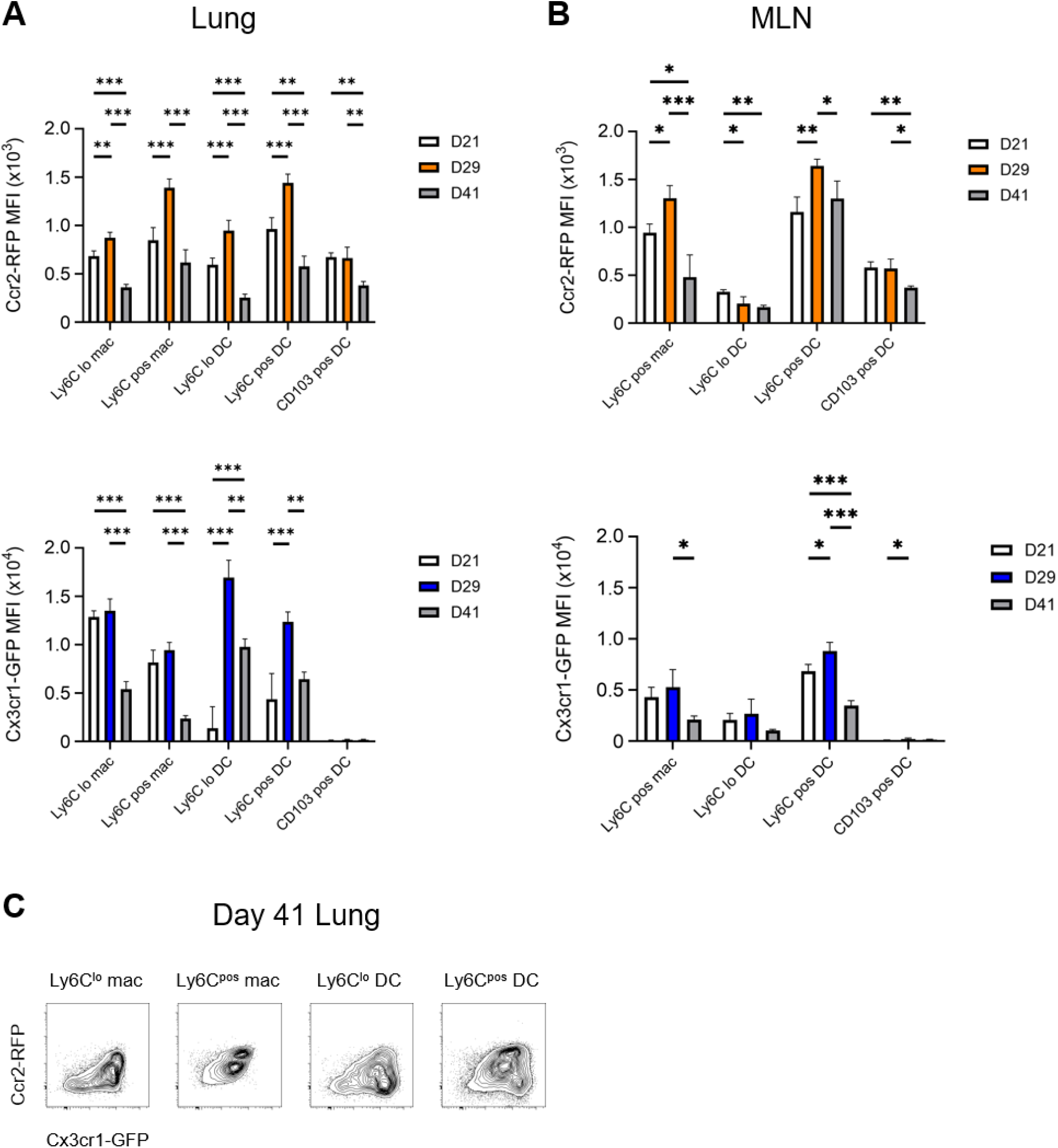
*Ccr2* and *Cx3cr1* expression in lung and MLN phagocytes over time. *Ccr2^RFP/+^*; *Cx3cr1^GFP/+^* mice were infected with Mtb for the indicated times, then lung (A) and MLN (B) cells were analyzed by flow cytometry. (C) Representative contour plots of the indicated populations from the lungs of mice infected for 41 days. All results presented as mean + SD and are derived from 1 experiment with 4 mice per genotype. Significance assessed by one-way ANOVA with multiple comparisons.

**Supplemental Figure 4:**
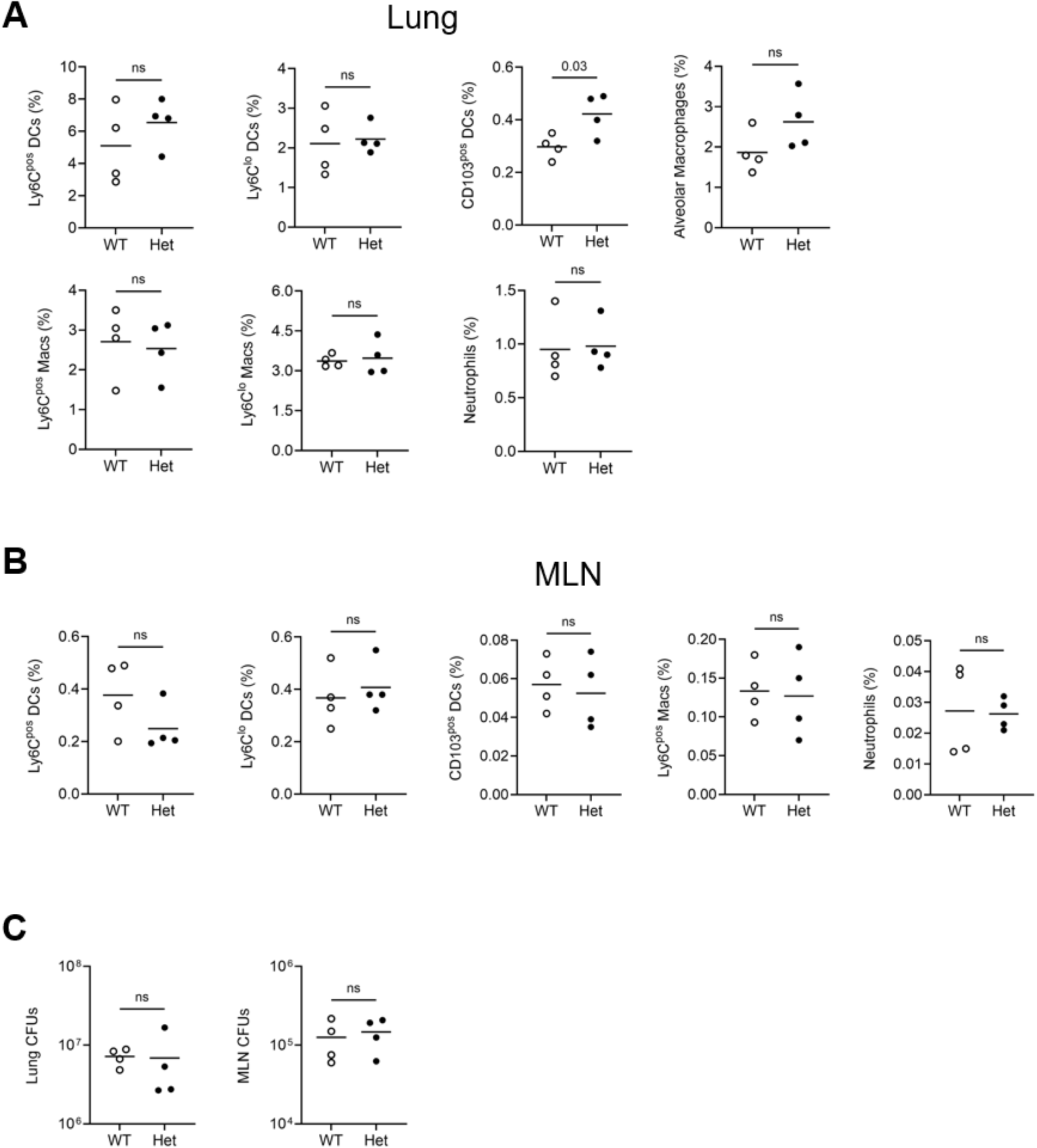
Comparison of Mtb-infected *Ccr2^+/+^*; *Cx3cr1^+/+^* and *Ccr2^RFP/+^*; *Cx3cr1^GFP/+^*mice. Mice were infected with Mtb for 4 weeks, then lungs and MLNs were harvested. (A-B) Frequencies of the indicated lung (A) and MLN (B) populations among live cells by flow cytometry. (C) Quantitation of Mtb CFUs in lungs and MLNs. Data are representative of 2 experiments with 4 mice per genotype. Horizontal bar represents the mean in all graphs. Significance assessed by *t* test.

**Supplemental Figure 5:**
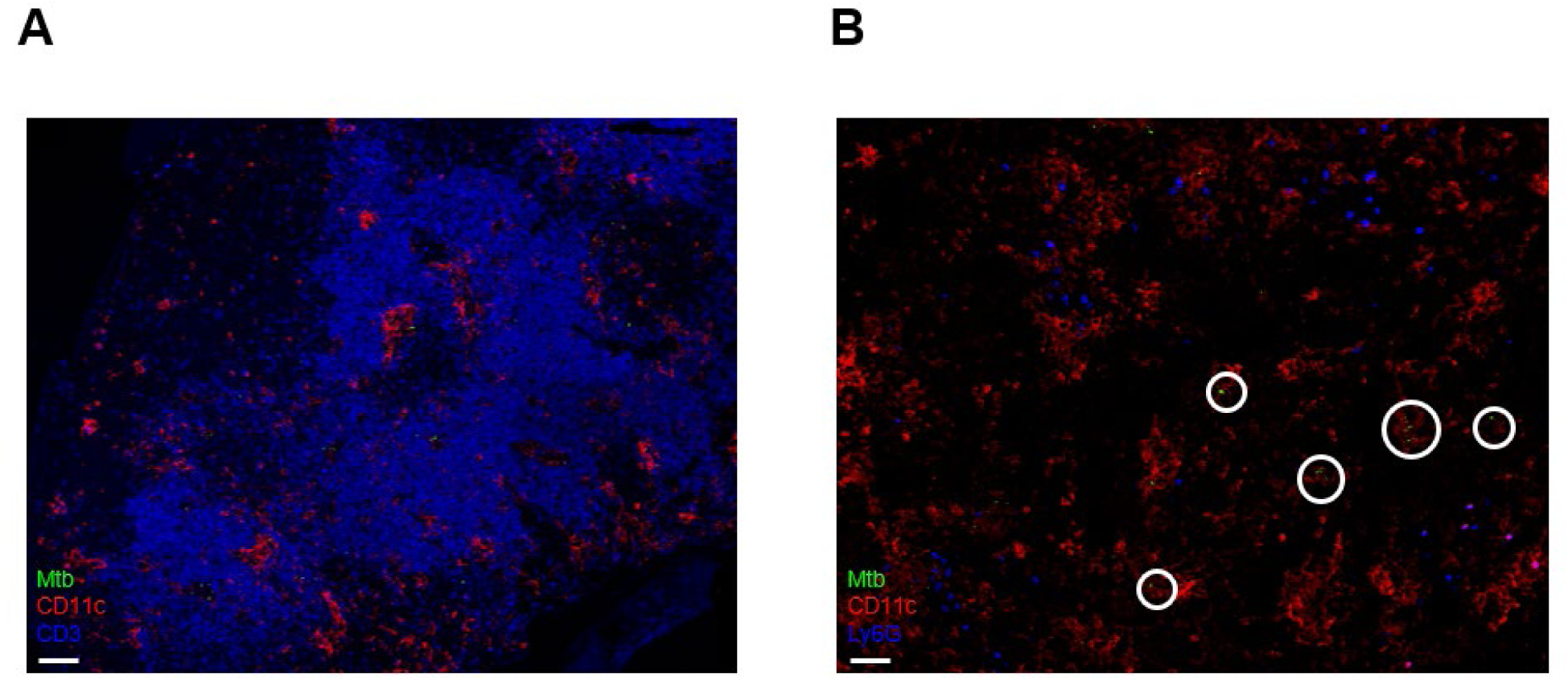
Positioning of neutrophils and T cells in MLNs of infected double knockout mice. DKO mice were infected for 4 weeks with YFP-expressing Mtb, then MLNs were sectioned and stained by immunofluorescence for CD11c and either CD3 (A) or Ly6G (B). Circles in (B) denote areas of CD11c and YFP co-localization without adjacent Ly6G staining. Images are representative of 2 experiments.

